# Getting Beyond Interruptive Alerts: Reducing Unintentional Duplicate Orders using a Visual Aid

**DOI:** 10.1101/640318

**Authors:** Steven Horng, Joshua W Joseph, Shelley Calder, Jennifer P Stevens, Ashley L O’Donoghue, Charles Safran, Larry A Nathanson, Evan L Leventhal

## Abstract

**Importance:** Electronic health records (EHRs) allow teams of clinicians to simultaneously care for patients but an unintended consequence could result in duplicate ordering of tests and medications.

**Objective:** We asked if a simple visual aid would reduce duplicate ordering of tests and medications for busy teams of clinicians in our emergency department by placing a red highlight around the checkbox of a computer-based order if previously ordered.

**Design:** We performed an interrupted time series to analyze all patient visits 1 year before and 1 year after the intervention. Significance testing was performed using a negative binomial regression with Newey-West standard errors, correcting for patient level variables and environmental variables that might be associated with duplicate orders.

**Setting:** The emergency department of an academic hospital in Boston, MA with 55,000 visits annually.

**Participants:** 184,722 consecutive emergency department patients.

**Exposure:** If an order had previously been placed during that ED visit, we cue the user by showing a red highlight around the checkbox of that order.

**Main Outcome:** Number of unintentional duplicate orders.

**Results:** After deployment of the non-interrupting nudge, the rate of unintentional duplicates for laboratory orders decreased 49% (incidence rate ratio 0.51, 95% CI 0.45-0.59) and for radiology orders decreased with an incidence rate ratio of 0.60 (0.44-0.82). There was no change in unintentional medication duplicate orders. We estimated that the nudge eliminated 17,936 clicks in our EHR.

**Conclusions and Relevance:** Passive visual queues that provide just-in-time decision support are effective, not disruptive of workflow, and may decrease alert fatigue in busy clinical environments.

**Key Points:** *Question:* Can a simple visual aid reduce duplicate ordering in an electronic health record?

*Findings:* In this interrupted time series, the rate of unintentional duplicates for laboratory orders decreased 49% and for radiology orders decreased 40%. There was no change in unintentional medication duplicate orders. We estimated that the nudge eliminated 17,936 clicks in our EHR.

*Meaning:* Quality improvement often relies on changing clinician behavior. We believe guiding clinicians to a right action is better than telling the clinician they have already made an error. Our approach will help reduce alert fatigue and lessen clinician complaints about EHRs.

## 1. BACKGROUND AND SIGNIFICANCE

Electronic health records (EHRs) could improve patient safety by facilitating communication, providing access to information, assisting with calculations, monitoring, and providing decision support.^12^ But the very presence of EHRs may lead to new unintended consequences, like increasing the likelihood that healthcare providers might be more likely to overlook existing orders and duplicate work.^7, 8^ Duplicate orders may be the intention of a provider looking to measure a lab value or radiology study serially. However, duplicate orders may also be markers of poor communication between two providers caring for the same patient, or may be an order placed on the wrong patient, a so-called “near miss”. ^9^

Common strategies to reduce duplicate orders include additional training for users, downstream workflow mitigation (such as screening by pharmacy, laboratory, or radiology), or interruptive alerts, all efforts to block healthcare providers from doing the wrong thing. ^12^ Though interruptive alerts can be effective, they generally occur after the provider has completed the ordering process, which can result in increased click-count, cognitive dissonance, frustration, and alert fatigue. ^13, 14^ Interruptive alerts can also disrupt current thought processes, which may lead to even more errors.^15, 16–19^

Larry Weed proposed another approach. He suggested that EHRs could both guide and teach. Passive just-in-time reminders could also be used to help guide users to the right thing, rather than preventing them from doing the wrong thing. The level of interruption to clinician workflows, whether guiding, preventing, or both, should be tailored to the severity and immediacy of the harm being prevented.^3^

The time sensitive nature of Emergency Department (ED) visits, combined with the need for parallel workflows and team-based care makes the ED setting particularly vulnerable to unintended duplicate orders. ^10, 11^ The goal of this investigation is to show the effect of a passive inline just-in-time decision support, functioning as a nudge for providers, on reducing unintentional duplicate orders in the emergency department.

## 2. METHODS AND METHODS

### 2.1 Study Design

We performed an interrupted time series to analyze all patient visits 1 year before and 1 year after each intervention.

### 2.2 Setting and Selection of Participants

The study was performed in a 55,000 visits/year Level I trauma center and tertiary, academic, adults-only, teaching hospital. All consecutive ED patient visits 1 year before and 1 year after each intervention were included in the study. No visits were excluded. The EHR used was custom developed at the institution.

### 2.3 Intervention

We implemented a user interface within our CPOE system in the ED that provides passive inline just-in-time duplicate order decision support while the user is in the process of placing an order. If an order had previously been placed during that ED visit, we cue the user by showing a red highlight around the checkbox of that order. (*Figure 1*) The intervention was performed as a phased implementation starting with laboratory orders (8/13/13), followed by medication orders (2/3/14), and lastly radiology orders (12/12/14).

**Figure 1:**
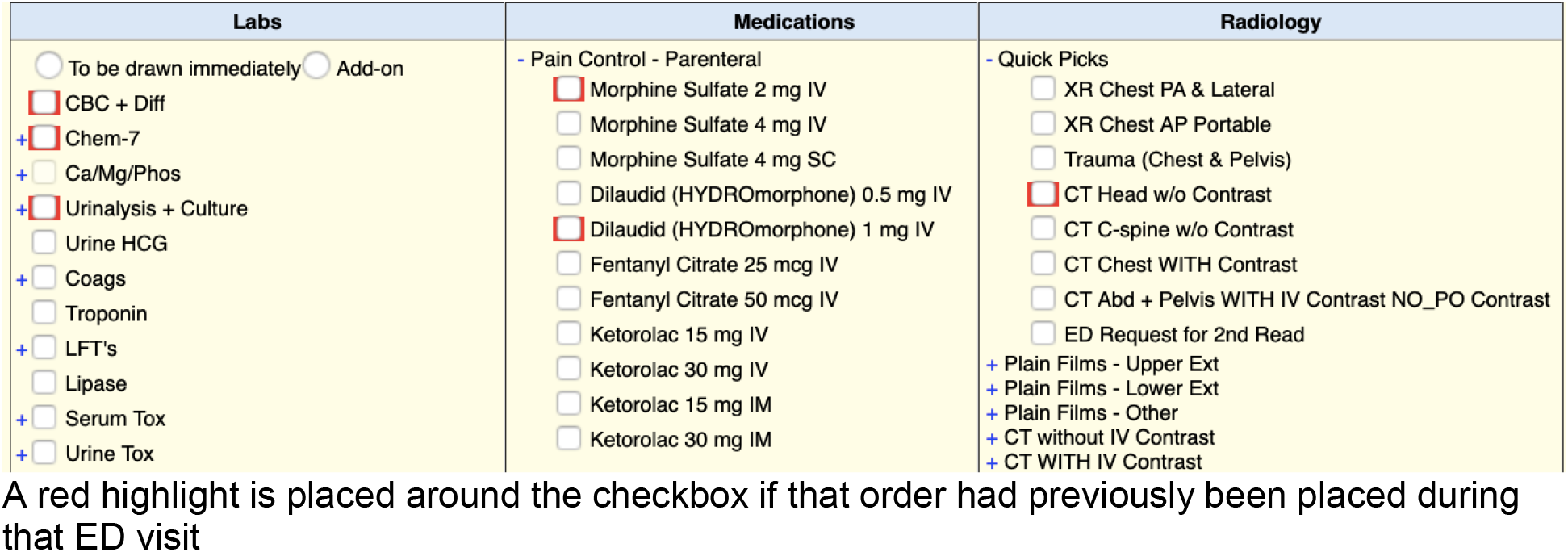
Passive inline just-in-time duplicate order decision support (red highlight)

### 2.4 Duplicate Order Algorithm

An order was determined to be a duplicate if two identical orders were placed, and then one was subsequently cancelled. For the purposes of this algorithm, a laboratory order is uniquely identified by an internal laboratory code, which is then mapped to a LOINC code. Medication orders are uniquely identified by the combination of an internal medication code, route, and dose. For example, two orders for morphine 5 mg IV once would be considered a duplicate order, while an order for morphine 5 mg IV once and morphine 2 mg IV once would not be considered a duplicate. Each internal medication code is then mapped to a medication entry in First Data Bank. Radiology orders are uniquely identified by a Simon-Leeming code.^24^

### 2.5 Methods of Measurement

Data was collected for each patient visit and then aggregated into 8 hours bins, corresponding to the 8 hour shifts worked by clinicians (7a-3p, 3p-11p, 11p-7a). For each bin, the mean patient age was calculated, as well as the percent of female patients, percent of patients with an emergency severity index (ESI) greater than or equal to 3, percent of patients with English as a primary language, percent discharged, and percent admitted. We also collected environmental variables to describe the state of the emergency department when the patient first arrived in the ED. These environmental variables included the current number of patients in the department, number of patients in observation, number of new patients, number of ICU bed requests, number of telemetry bed requests, number of floor bed requests, and number of boarders. We then calculate the mean for each of these environmental variables for each bin. ESI is on a scale from 1 to 5, with 1 being critically ill patients requiring immediate intervention and 5 being the least acute. Boarders are patients who have been admitted to the hospital, but have not yet received a bed after 2 hours.

### 2.6 Outcome Measure

The outcome measure was the number of unintentional duplicate orders in each bin for each of the order types: laboratory, medication, and radiology.

### 2.7 Data Analysis

Data was analyzed using an interrupted time series analysis. Since we performed a phased implementation, we shifted the days so that the intervention occurred on the same day for each order type. Orders were aggregated into 8 hour bins.

We performed individual negative binomial models for each of the duplicate order types: laboratory orders, medications, and radiology studies. The total number of orders, less duplicates, were included as an offset term. All models were controlled at the shift-level, including average patient age, gender composition of patients, whether patients were native English speakers, percent of patients with an emergency severity index 3 or greater, percent of patients discharged home, percent of patients admitted, number of patients in the department, number of patients in the waiting room, number of observation patients, number of new patients, number of ICU beds requested, number of telemetry beds requested, and number of boarders. Fixed effects in the model were included for time of day at the shift level, day-of-week, and month-of-sample, as duplicate orders may follow a cyclical pattern throughout the day, week, and year, respectively. We report both the immediate effect, or the level change, of the intervention on duplicate orders and an estimate of the change in trend of duplicate orders after the intervention. Coefficients were transformed into incidence rate ratios (IRRs) in Table 2. Stata/SE 14.2 software was used for statistical analysis. A more detailed description of the statistical analysis can be found in the Appendix.

### 2.7 Human Subjects Research Statement

A determination was made by our Committee on Clinical Investigation that this did not constitute human subjects research and was considered exempt from review. (#2019D000239)

## 3. RESULTS

### 3.1 Summary Statistics

A total of 730 days were analyzed. Patient demographics and order summary statistics are reported in *Table 1*.

**Table 1:** Patient Demographics and Order Summary Statistics.

|  | Laboratory<br>Mean (SD) |  |  | Medications<br>Mean (SD) |  |  | Radiology<br>Mean (SD) |  |  |
| --- | --- | --- | --- | --- | --- | --- | --- | --- | --- |
|  | <u>Before</u> | <u>After</u> | <u>p-value</u> | <u>Before</u> | <u>After</u> | <u>p-value</u> | <u>Before</u> | <u>After</u> | <u>p-value</u> |
| **Patient Age** | **50.32**<br>(4.64) | **51.01**<br>(4.29) | **<0.001** | **50.55**<br>(4.58) | **51.15**<br>(4.38) | **0.002** | 51.09<br>(4.39) | 51.37<br>(4.05) | 0.120 |
| **% Female Patients** | 53.52<br>(8.40) | 53.70<br>(8.16) | 0.604 | 53.36<br>(8.36) | 53.75<br>(8.27) | 0.273 | 53.90<br>(8.17) | 53.46<br>(8.30) | 0.213 |
| **% Patients with ESI >= 3** | **62.26**<br>(9.51) | **61.23**<br>(9.42) | **0.010** | **62.03**<br>(9.75) | **60.29**<br>(9.65) | **0.000** | **60.63**<br>(9.32) | **58.59**<br>(10.09) | **0.000** |
| **% Patients with English as primary language** | 96.54<br>(3.4) | 96.38<br>(3.53) | 0.455 | 96.45<br>(3.42) | 96.25<br>(3.54) | 0.168 | 96.23<br>(3.55) | 96.50<br>(3.33) | 0.062 |
| **% Discharged** | 59.23<br>(9.04) | 59.95<br>(8.3) | 0.455 | **59.11**<br>(8.51) | **58.02**<br>(8.73) | **0.003** | 58.30<br>(8.56) | 57.70<br>(8.74) | 0.105 |
| **% Admitted** | 36.56<br>(8.58) | 36.53<br>(7.93) | 0.934 | 36.73<br>(8.21) | 37.14<br>(8.38) | 0.243 | 36.91<br>(8.06) | 37.12<br>(8.12) | 0.546 |
| **No. Patients in Department** | **39.1**<br>(8.75) | **40.2**<br>(8.85) | **0.003** | **38.85**<br>(8.71) | **42.31**<br>(8.98) | **0.000** | **41.38**<br>(9.04) | **45.61**<br>(8.94) | **0.000** |
| **No. Patients in Waitroom** | 3.66<br>(3.60) | 3.95<br>(3.92) | 0.078 | **3.64**<br>(3.65) | **4.31**<br>(4.10) | **0.000** | **4.12**<br>(4.00) | **5.64**<br>(4.80) | **0.000** |
| **No. Patients in Observation** | **9.47**<br>(3.69) | **8.80**<br>(3.61) | **0.000** | **8.98**<br>(3.62) | **9.4**<br>(3.71) | **0.006** | **9.24**<br>(3.73) | **10.46**<br>(3.91) | **0.000** |
| **No. New Patients** | 6.86<br>(2.62) | 6.81<br>(2.57) | 0.646 | 6.77<br>(2.59) | 6.90<br>(2.56) | 0.233 | 6.86<br>(2.55) | 6.97<br>(2.56) | 0.314 |
| **No. ICU Bed Requests** | **0.84**<br>(0.61) | **0.91**<br>(0.67) | **0.018** | **0.86**<br>(0.63) | **1.01**<br>(0.74) | **0.000** | **0.94**<br>(0.69) | **1.26**<br>(0.87) | **0.000** |
| **No. Telemetry Bed Requests** | **2.3**<br>(1.47) | **2.67**<br>(1.74) | **0.000** | **2.40**<br>(1.51) | **3.12**<br>(2.01) | **0.000** | **2.82**<br>(1.83) | **3.8**<br>(2.17) | **0.000** |
| **No. Floor Bed Requests** | 3.53<br>(1.88) | 3.64<br>(1.95) | 0.178 | **3.41**<br>(1.82) | **3.96**<br>(2.09) | **0.000** | **3.81**<br>(2.03) | **4.71**<br>(2.44) | **0.000** |
| **No. Boarders** | **2.48**<br>(1.88) | **1.34**<br>(1.67) | **0.000** | 1.79<br>(1.83) | 1.85<br>(2.07) | 0.448 | **1.50**<br>(1.79) | **2.98**<br>(2.64) | **0.000** |
| **No. of Orders** | 306.51<br>(149.31) | 308.69<br>(141.95) | 0.725 | **140.82**<br>(64.12) | **155.77**<br>(71.84) | **0.000** | **56.93**<br>(26.76) | **60.52**<br>(28.05) | **0.002** |
| **No. of**<br>**Unintentional**<br>**Duplicate Orders** | **4.1**<br>**(5.18)** | **2.49**<br>**(3.39)** | **0.000** | **0.21**<br>**(0.51)** | **0.26**<br>**(0.55)** | **0.016** | **0.87**<br>**(1.18)** | **0.71**<br>**(0.98)** | **0.001** |
| **% Duplicate**<br>**Orders** | **1.29**<br>**(1.65)** | **0.81**<br>**(1.24)** | **0.000** | **0.13**<br>**(0.34)** | **0.17**<br>**(0.42)** | **0.011** | **1.59**<br>**(2.38)** | **1.21**<br>**(1.98)** | **0.000** |

**Table 2:**
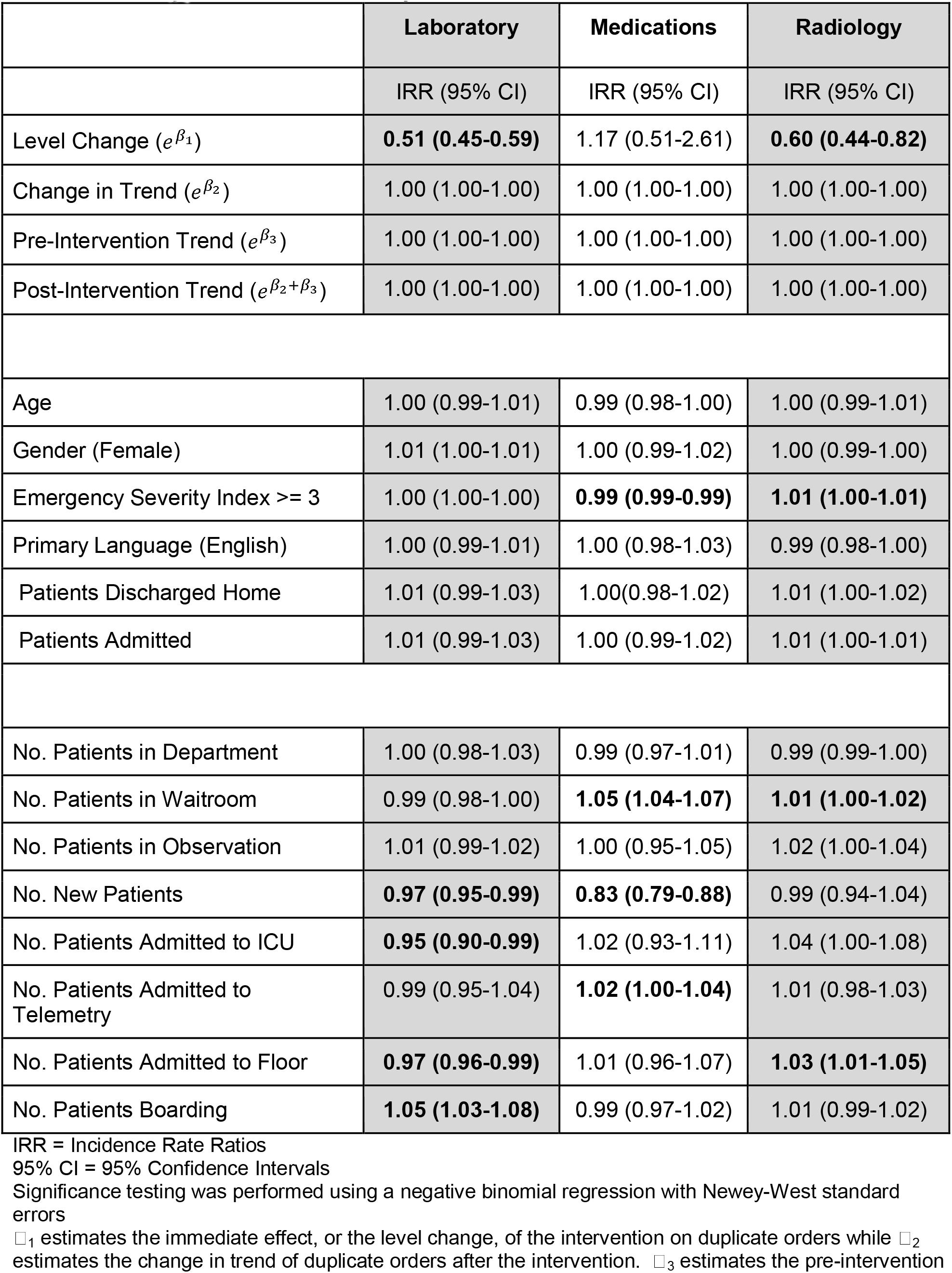
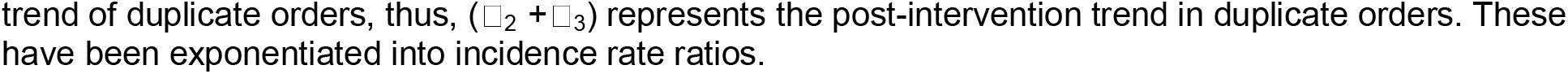
Interrupted Time Series Analysis.

### 3.2 Main Results

After the intervention, there was a significant level change with a decrease in laboratory duplicate orders incidence rate ratio (IRR) 0.51 (95% confidence interval (CI) 0.45-0.59) and radiology duplicate orders IRR 0.60 (0.44-0.82). Additionally, while the estimate for medication duplicate orders is positive, it was not statistically significant. There was no change in trend for any of the intervention groups. We report these results from the interrupted time series in *Table 2*.

For patient variables, an ESI >= 3 (lower acuity) was also associated with more radiology duplicate orders after the intervention IRR 1.01 (1.00-1.01), but fewer medication duplicate orders, IRR 0.99 (0.99-0.99).

For environmental variables, the number of patients in the waitroom was associated with an increase in medication and radiology duplicate orders. The number of new patients during that shift was associated with a decrease in duplicate orders in the laboratory and medication intervention groups. The number of patients admitted to the ICU during that shift was associated with a decrease in the number of laboratory duplicate orders. Number of patients admitted to telemetry was associated with an increase in medication duplicate orders. Number of patients admitted to the floor during that shift was associated with a decrease in laboratory duplicate orders, while associated with an increase in radiology duplicate orders. Lastly, the number of patients boarding during that shift was associated with an increase in laboratory duplicate orders.

We also report the top 5 duplicate orders for each order type in *Supplemental Table* to better illustrate the most common unintentional duplicate orders.

## 4. DISCUSSION

### 4.1 Just-in-time Passive Decision Support

In our emergency department, a red highlight around an order’s checkbox (Figure 1) significantly reduced duplicate orders for both laboratory testing and radiological procedures but not medications. We have demonstrated a simple, yet effective method of reducing duplicate laboratory and radiology orders that avoids many of the downsides of traditional, interruptive methods. This method was evaluated in a busy clinical setting that relies on effective and efficient team work to care for patients. It takes 9 clicks and the typing of a password to cancel an order in our EHR, which takes 30 seconds to complete. The predicted reduction in unintended duplicate orders saved 17,936 clicks in the year after the intervention, which amounts to 16 hours and 36 minutes of regained productivity.

We took a user-centered approach to user interface design and followed the best practices of clinical decision support to minimize interruptions unless they are clinically necessary.^3^ We placed the decision support for duplicate orders just-in-time, which is at exactly the time the decision to order was being made, rather than an interruptive alert, which is after a decision has already been made. We furthermore placed our decision support inline, so that the decision support was in the same area of focus as the intended action.

At baseline, we had relatively few medication duplicate orders compared to laboratory and radiology duplicate orders, which is consistent with ED workflow. Nurses often order laboratory and radiographic studies per clinical protocol, but do not routinely order medications, except in an emergency when a physician gives a nurse a verbal order. A medication duplicate order would normally only occur if two physicians, for example, a resident and attending, both ordered the same medication.

### 4.2 Comparison to Interruptive Alerts

Interruptive alerts fundamentally make the wrong thing hard to do, but often with substantial cost to physician time. By contrast to our strategy, Procop et al used a hard stop phone call requiring physicians to call the laboratory to order a duplicate from a list of more than 1,200 lab tests.^25^ This intervention was successful at reducing duplicate lab orders, with only 3% of interruptive alerts being overridden by a phone call, resulting in a cost avoidance of $183,586 over 2 years accounting for materials and laboratory personnel labor, but failing to account for the physician labor used to initiate the call.Though poor usability can be a powerful deterrent, we believe good usability is a more sustainable approach that respects human autonomy and promotes user satisfaction and physician longevity.

Interruptive alerts can also lead to patient harm. In 2006, a multi-center randomized controlled trial was performed to evaluate an interruptive alert to reduce concomitant orders for warfarin and trimethoprim-sulfamethoxazole.^26^ Although, the interruptive alert was successful in reducing this drug pair ordering with an adjusted odds ratio of 0.12 (95% CI 0.045-0.33), it also had the unintended consequence of 4 patients who had delays in treatment. The harm was believed sufficient that the IRB terminated the study early.

### 4.3 Intervention User Interface Design

We took a user-centered approach to user interface design and followed the best practices of clinical decision support to minimize interruptions unless they are clinically necessary.^3^ We placed the decision support for duplicate orders just-in-time, which is at exactly the time the decision to order was being made, rather than an interruptive alert, which is after a decision has already been made. We furthermore placed our decision support inline, so that the decision support was in the same area of focus as the intended action. A common alternative to providing just-in-time decision support would be to display a list of current orders for a patient on the order entry screen. A user would first need to commit these orders to short term memory and then apply them to the order entry screen. As patients can have tens of orders, this can quickly overwhelm a user’s working memory, leading to unintended cognitive errors.

Rather than leverage the user’s computational ability to do duplicate order checking, allergy checking, or drug-drug interaction checking, we use a computer’s computational ability. Unlike a human, a computer’s computational ability is highly reliable, fast, and does not fatigue. We provide this computational capacity in real-time at the time of screen draw with sub-second response times. Though in the past, computational resources were limited and such computations can not be performed in real-time and needed to be precalculated, advances in computational capacity has eliminated this concern.

### 4.4 Additional Use Cases

We also use these same techniques for duplicate order reminders in our order sets, transforming the order set itself into a visual checklist, as shown in *Figure 3*. These visual checklists engage the visual and spatial reasoning portions of the brain, offloading the language centers. These same techniques can also be used to guide users when using a clinical decision rules, as shown in *Supplemental Figure 1*, suggesting the correct responses based on automated information retrieval. In the same way, such methods can also be used to integrate predictive machine learning algorithms into clinical workflows by guiding users to the correct response along with some explanation from the algorithm to show its work. Since a human is still in the loop, this method would be tolerant to less than perfect performance metrics as newer machine learning models are being created.

Though for this application we employed passive inline JIT decision support alone, this approach need not be mutually exclusive with interruptive alerts. For example, we have also implemented common drug decision support in CPOE and prescribing modules for allergies, pregnancy, and drug-drug interactions using both approaches. Allergy decision support for CPOE and prescribing modules is very amenable to this approach, as some patients can have multiple drug allergies, shown in *Supplemental Figure 2*. Though EHRs commonly provide JIT decision support by enumerating a patient’s allergies on the order screen, users must still commit these allergies to working memory, and then apply them by drug class to each potential order. This workflow creates multiple failure modes: the committing of allergies to working memory, the grouping of allergies to drug classes, and finally the cross-referencing of drug classes to orders. Our inline decision support offloads this cognitive process from the user, freeing their attention to focus on other clinical tasks.

**Figure 2:**
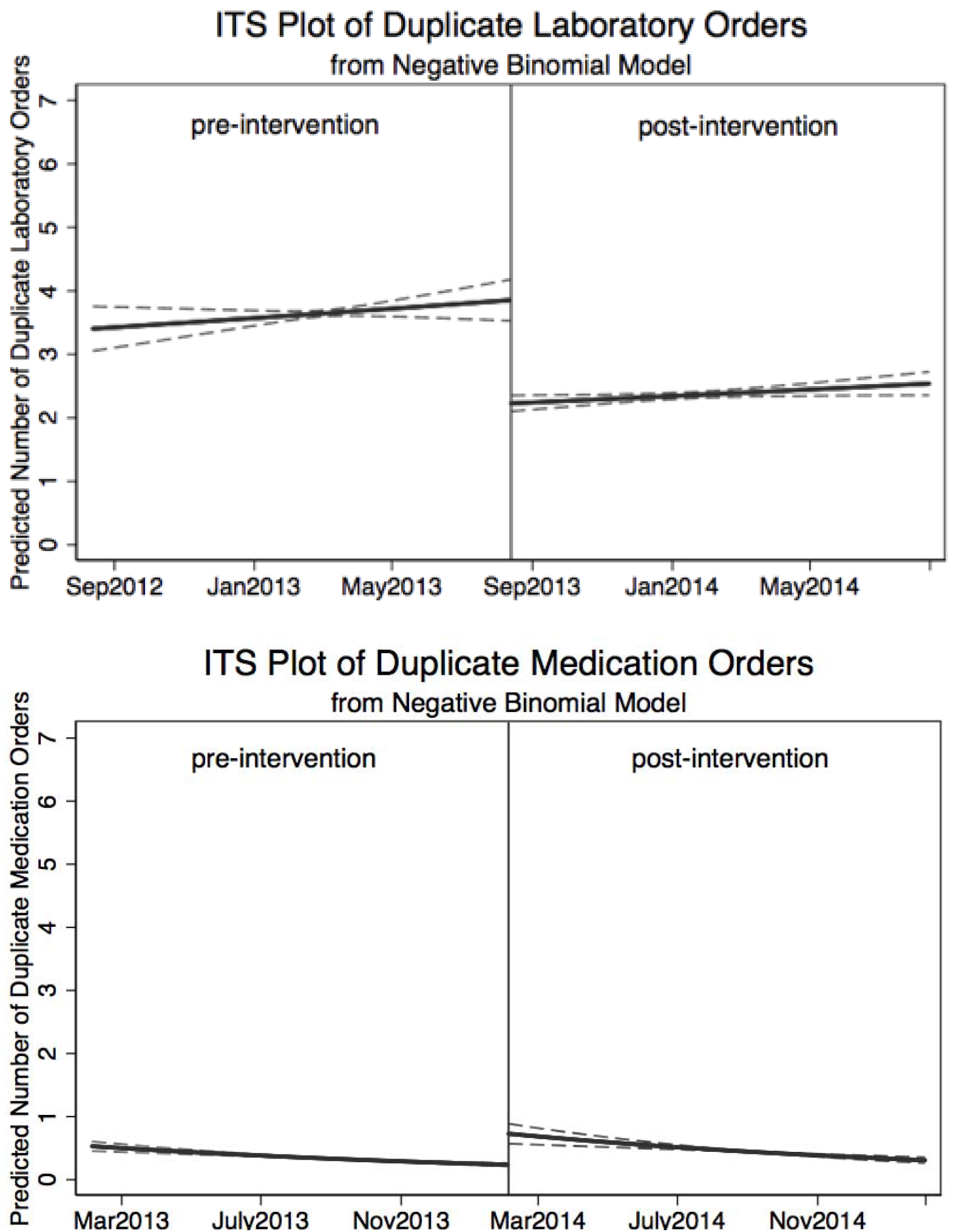
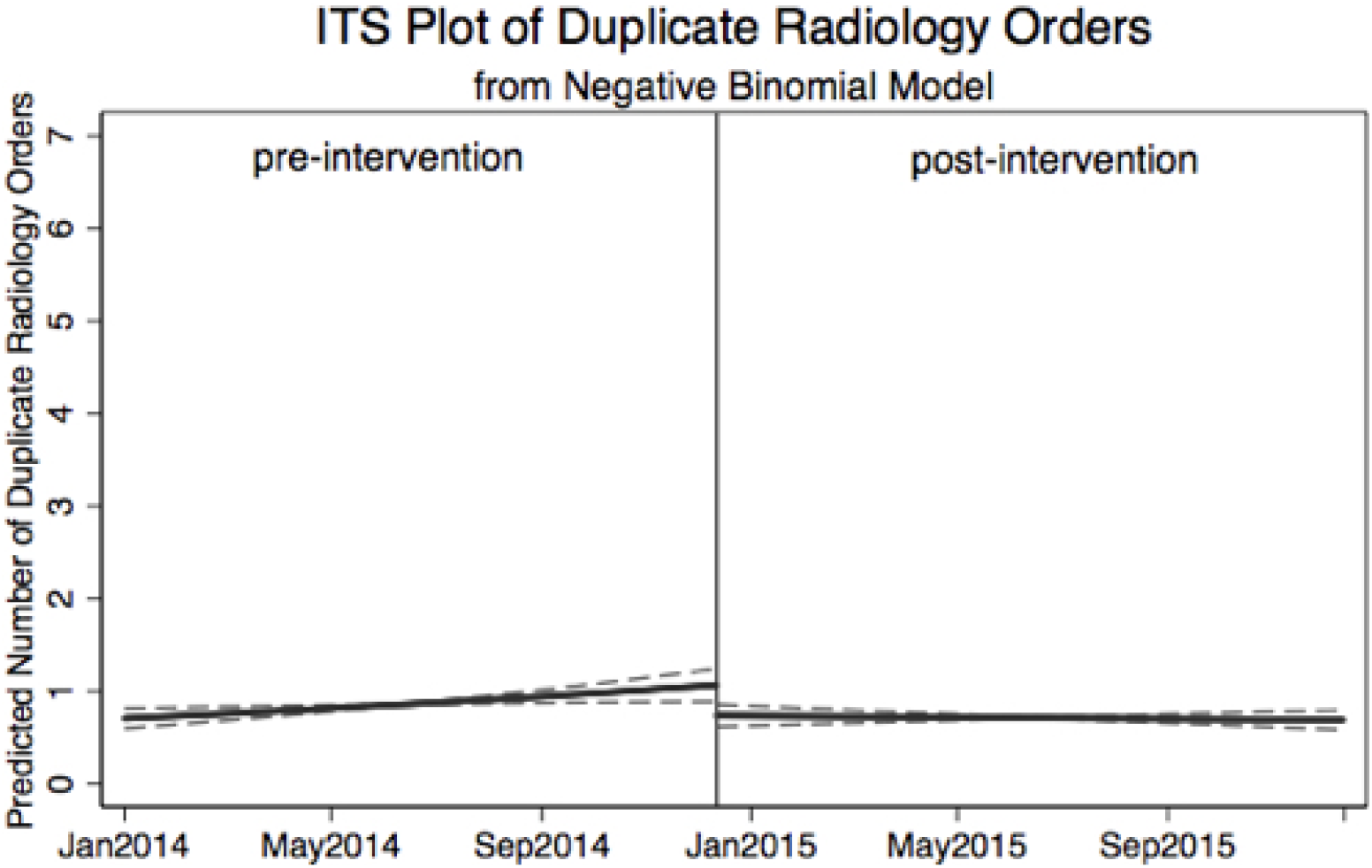
Interrupted Time Series of Unintentional Duplicate Orders.

**Figure 3:**
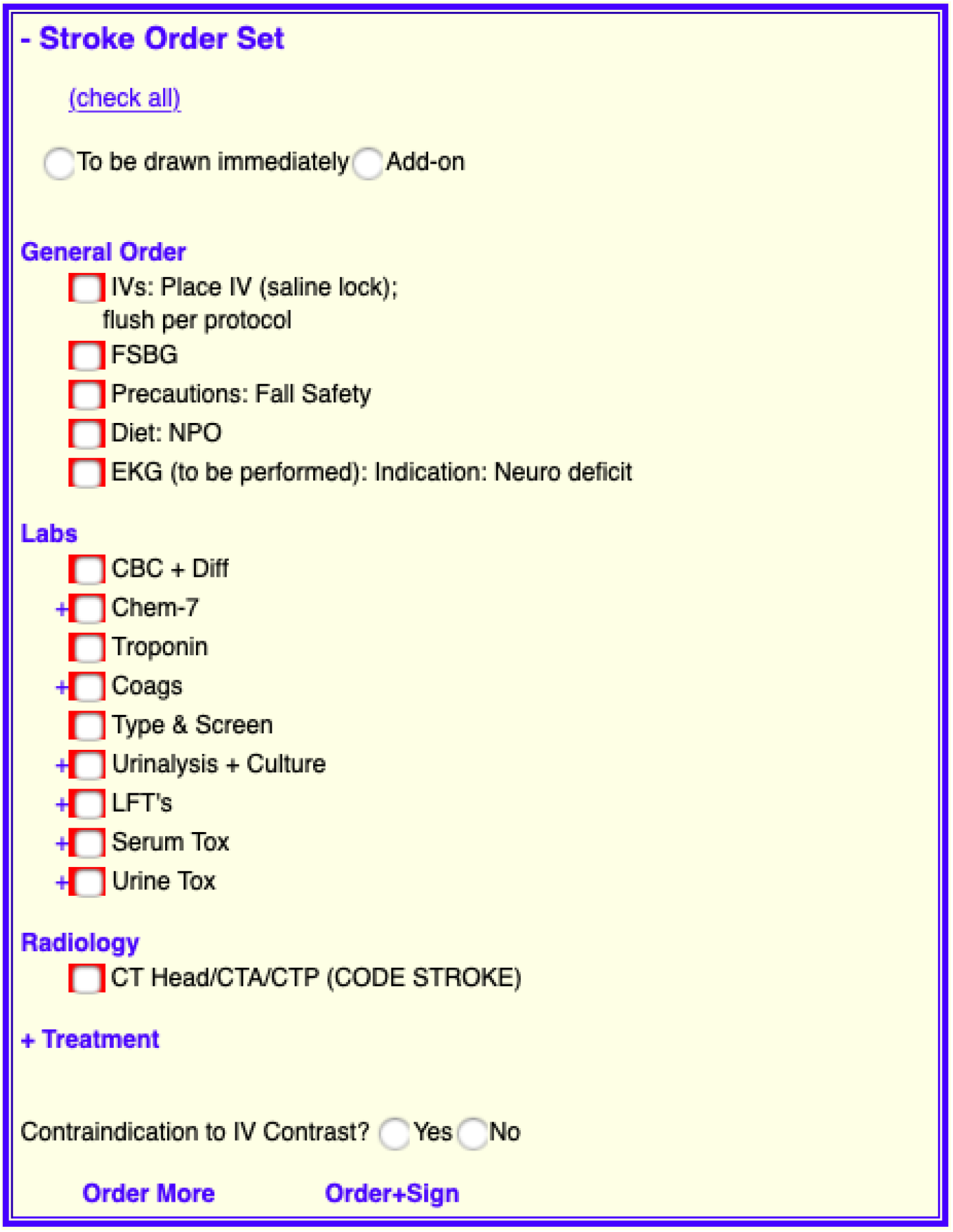
Order Sets as Visual Checklists.

### 4.5 Limitations

First, we conducted an observational study, and our results may be the consequence of unmeasured confounding variables. Although randomized controlled trials remain the gold-standard for assessing the effect of interventions, it is not always possible in informatics-based interventions in the ED. To mitigate the risks of observation design, we performed an interrupted time series analysis and corrected for every patient level variable or environmental variable collected in our EHR that might be associated with duplicate orders. Further, in our intervention, the effect was unlikely due to chance or secular trends, as the individual interventions on laboratory, medications, and radiology orders were phased in at different times and then time shifted so that the intervention date occurred on the same date.

Second, our outcome measure only captured duplicate orders that were subsequently cancelled, using the act of cancellation as a surrogate for user intent. There were also unintentional duplicate orders that were never cancelled and subsequently carried out.

Third, our intervention and outcome measure only addressed exact duplicate orders. Semantic duplicate orders such as therapeutic duplication are also important and remain as future work. For example, a patient who is ordered acetaminophen/oxycodone should receive passive JIT decision support no only for acetaminophen/oxycodone, but also for its constituent ingredients: acetaminophen and oxycodone.

Finally, our study was also performed at an urban, academic, adults-only teaching hospital with a custom EHR. The effectiveness of this intervention may differ in other care settings with different baseline unintentional duplicate orders and mitigation workflows.

### 4.6 Future Directions

Our implementation was specific to the ED, but should work in any care setting where clinical care teams must collaborate. Other care settings that have clinicians who are geographically dispersed would amplify the effect of such an intervention. Different care settings will require different thresholds for determining if an order is a duplicate. For example, in the ICU, that threshold might be 4-12 hours. In the hospitalist setting, that threshold may be 24 hours, while in the outpatient setting, it may be 1 week or 1 month. In the ED, having a simple binary representation of whether an order was a duplicate was intuitive and clinically meaningful. However, in the ICU, having multiple stages corresponding with last order time with additional information such as ordered 5 mins ago or ordered 6 hours ago might be important. We will iteratively develop new user interfaces given this increased complexity as we deploy this intervention across these different care settings.

## 5. CONCLUSION

Passive just-in-time decision support is an effective alternative to interruptive post-order interruptive alerts for reducing duplicate order entry. We believe guiding clinicians to a right action is better than telling the clinician they have made an error. Our approach will help reduce alert fatigue and lessen clinician complaints about EHRs.

## APPENDIX

We modeled duplicate orders for each of the three order types with the following negative binomial regression:

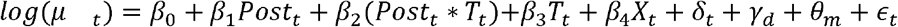

where *μ* _*t*_ represents the expected number of duplicate orders of its type. The total number of orders, less duplicates, were included as an offset term. *Post*_*t*_ is an indicator variable that equals 1 after the intervention date for that order type and 0 before. *T*_*t*_ is the number of shifts at time *t* since the beginning of the sample for that order type, where the sample begins 1 year prior to the intervention date. *X*_*t*_ is a vector of controls at time *t* at the shift-level, including average patient age, gender composition of patients, whether patients were native English speakers, percent of patients with an emergency severity index 3 or greater, percent of patients discharged, percent of patients admitted, number of patients in the department, number of patients in the waiting room, number of observation patients, number of new patients, number of ICU beds requested, number of telemetry beds requested, and number of boarders. *δ*_*t*_,*γ*_*d*_, and *θ*_*m*_ are shift fixed effects, day-of-week fixed effects, and month-of-sample fixed effects that control for how duplicate orders may follow a cyclical pattern throughout the day, week, and year, respectively. *ϵ*_*t*_ are Newey-West standard errors, which are robust to autocorrelation and heteroskedasticity. All tests used a significance level (α) of 0.05.

The coefficient of interest are *β*_1_ and *β*_2_. *β*_1_ estimates the immediate effect, or the level change, of the intervention on duplicate orders while *β*_2_ estimates the change in trend of duplicate orders after the intervention. *β*_3_ estimates the pre-intervention trend of duplicate orders, thus, (*β*_2_ + *β*_3_)represents the post-intervention trend in duplicate orders. Coefficients are transformed into incidence rate ratios (IRRs) in Table 2, such that 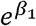 estimates the immediate increase in the incidence of duplicate orders after implementation, 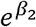 estimates the gradual change in the incidence of duplicate orders over time after implementation, 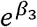 estimates the pre-intervention trend in incidence over time, and 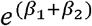 represents the post-intervention trend in incidence of duplicate orders.

We initially used a Poisson regression to model counts. However, a likelihood ratio test of alpha = 0 generated a large test statistic of 2734.83 with an associated p-value of < 0.001. This suggests that our duplicate order response variables are over-dispersed and therefore we used the more general negative binomial model.

Because we estimated three separate models for each of the types of duplicate orders, the coefficients on the control variables are estimated separately. One may be concerned that the effect of these control variables should not be different across the three models. For instance, the impact of average age on the probability of duplicate orders should likely be the same across the three types of orders as age should not impact the probability of a laboratory duplicate order differently than a medication or radiology duplicate order. Indeed, when testing the equality of the coefficients on the controls across the three models, we find that they are not statistically different from each other, with the only exception being number of patients in the waitroom. Thus, we re-estimated our models with the imposed constraint that the estimated coefficients on the control variables that were not statistically different from each other are equal across models. This allows us to estimate the impact of the intervention on each of the order types while jointly estimating the coefficients on the controls overall. In this alternative specification, our estimates of the effect of the program do not change in magnitude, direction, or significance.

Stata/SE 14.2 software was used for statistical analysis.

---

**Supplemental Table:** Top 5 Unintentional Duplicate Orders.

|  | **Laboratory Duplicate Orders** |  |  |  |
| --- | --- | --- | --- | --- |
|  | **<u>BEFORE</u>** |  | **<u>AFTER</u>** |  |
|  | Rank | Count (%)* | Rank | Count (%) * |
| Chem-7 | 1 | 726 (16%) | 1 | 361 (13%) |
| CBC/DIF | 2 | 615 (13%) | 2 | 269 (10%) |
| Coags | 3 | 239 (5%) | 4 | 171 (6%) |
| Urinalysis | 4 | 197 (4%) | 11 | 76 (3%) |
| Lactate | 5 | 194 (4%) | 6 | 124 (5%) |
|  | ... |  | ... |  |
|  | **Total** | **4560** | **Total** | **2729** |
|  | **Medication Duplicate Orders** |  |  |  |
|  | **<u>BEFORE</u>** |  | **<u>AFTER</u>** |  |
|  | Rank | Count (%)* | Rank | Count (%) * |
| Hydromorphone (IV) | 1 | 53 (14%) | 1 | 58 (12%) |
| Diazepam (PO) | 2 | 33 (9%) | 2 | 42 (8%) |
| Morphine (IV) | 3 | 26 (7%) | 3 | 29 (6%) |
| Lorazepam (IV) | 4 | 11 (3%) | 9 | 11 (2%) |
| Aspirin (PO) | 5 | 8 (2%) | 33 | 3 (1%) |
|  | ... |  | ... |  |
|  | **Total** | **380** | **Total** | **503** |
|  | **Radiology Duplicate Orders** |  |  |  |

|  | <u>BEFORE</u> |  | <u>AFTER</u> |  |
| --- | --- | --- | --- | --- |
|  | Rank | Count (%)<sup>*</sup> | Rank | Count (%)<sup>*</sup> |
| Chest X-Ray (PA & Lat) | 1 | 149 (15%) | 2 | 101 (13%) |
| CT Head w/o Contrast | 2 | 118 (12%) | 3 | 86 (11%) |
| CT Abdomen & Pelvis w/ Contrast | 3 | 113 (12%) | 1 | 117 (15%) |
| CT C-Spine w/o Contrast | 4 | 45 (5%) | 7 | 26 (3%) |
| Chest X-Ray (Portable) | 5 | 43 (4%) | 4 | 34 (4%) |
|  | ... |  | ... |  |
|  | **Total** | **962** | **Total** | **791** |

<sup>\*</sup> Count (%) should only be compared vertically, not horizontally, as count (%) are not normalized for differences in the number of patients and total orders in each period. Please use the ranks to compare horizontally. Please refer to *Table 1* and *Table 2* for normalized results to compare between periods.

**Supplemental Figure 1:**
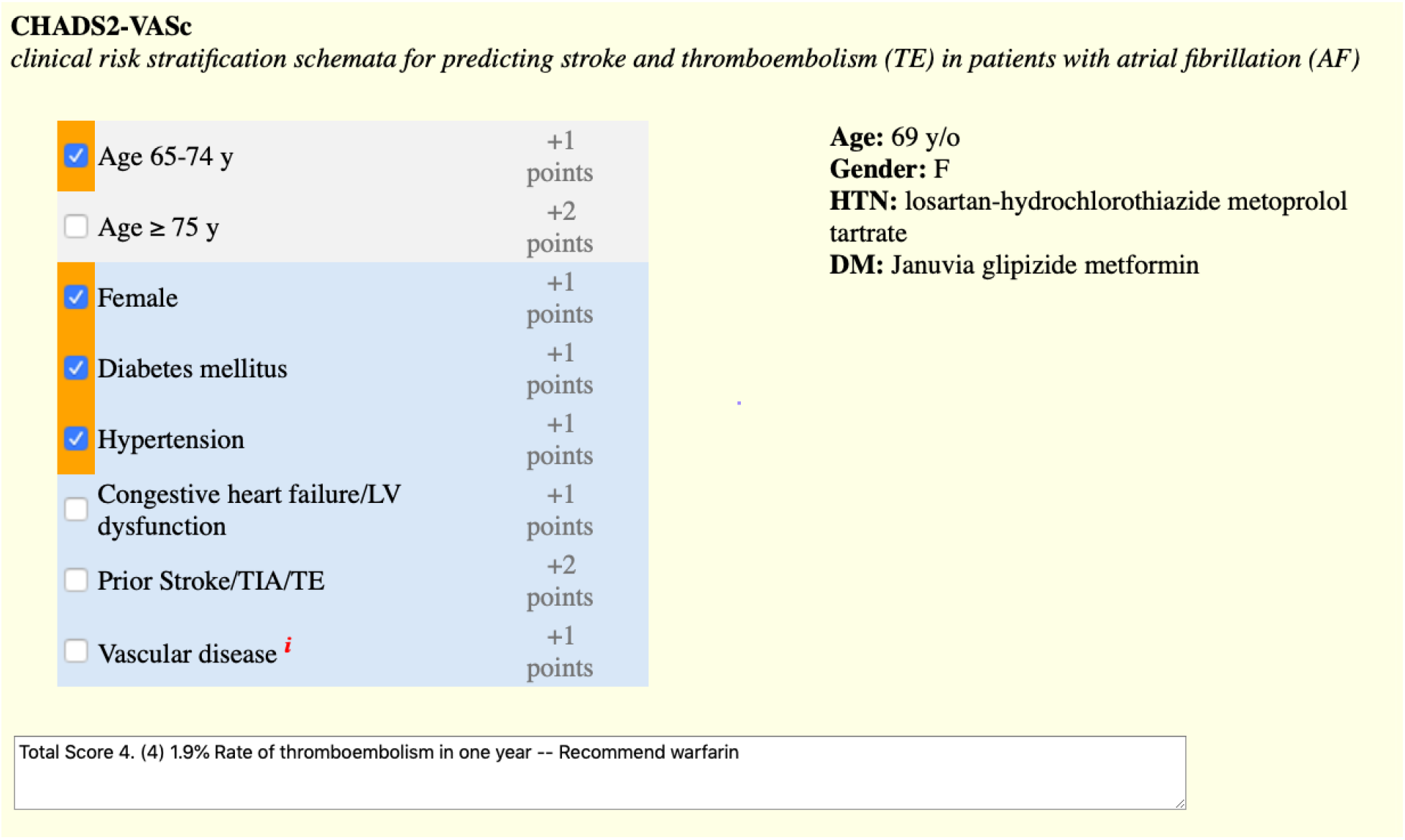
Suggestions for Clinical Decision Rules.

**Supplemental Figure 2:**
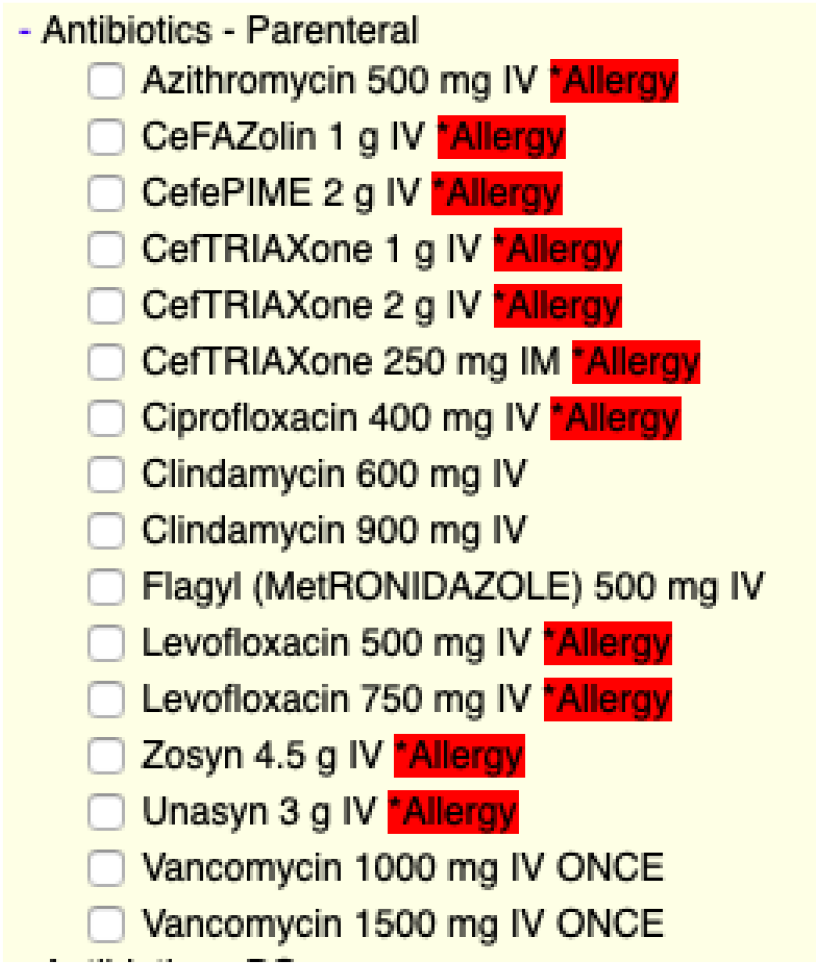
Allergy Decision Support.

